# Ecologically Significant Genetic Loci of *P. allii*: Contributions to Pathogenesis and Competition

**DOI:** 10.1101/2025.04.23.650278

**Authors:** Gi Yoon Shin, Stefanie De Armas, Guillermo A. Galván, María I. Siri, Mariah Rojas, Boris A. Vinatzer, Jo Ann E. Asselin, Paul Stodghill, Mei Zhao, Bhabesh Dutta, James Tambong, Brian H. Kvitko

**Affiliations:** Department of Plant Pathology, University of California Davis, Davis, CA, USA; Laboratorio de Microbiología Molecular, Montevideo, Facultad de Química, Universidad de la República, Uruguay; Departamento de Producción Vegetal (CRS), Facultad de Agronomía, Universidad de la República, Uruguay; School of Plant and Environmental Sciences, Virginia Tech, Blacksburg, VA, USA; Emerging Pests and Pathogens Research Unit, Robert W. Holley Center for Agriculture and Health, Agricultural Research Service, United States Department of Agriculture, Ithaca, NY, USA; Plant Pathology & Plant-Microbe Biology Section, School of Integrated Plant Science, Cornell University, Ithaca, NY, U.S.A; Department of Plant Pathology, College of Plant Protection, China Agricultural University, Bejing, R. T. China; Department of Plant Pathology, University of Georgia, Tifton, GA, USA; Ottawa Research and Development Centre, Agriculture and Agri-Food Canada, Ottawa, ON, Canada; Department of Plant Pathology, University of Georgia, Athens, GA, USA

**Author notes:** **Corresponding author** Brian H. Kvitko.

**Keywords:** *Pantoea allii*, pangenomics, HiVir, Halophos, Type 6 Secretion System, Tailocin, Ice nucleation

## Abstract

*Pantoea allii*, one of four *Pantoea* species known to cause onion center rot, is infrequently isolated from onion compared to its closely related onion-pathogenic sister taxa. To better understand the genomic diversity and genetic determinants of pathogenicity in this species, we analyzed a collection of 38 *P. allii* strains isolated from two primary ecological niches, plants and water, across three continents using comparative genomics and phylogenetic approaches. Core-genome phylogeny, average nucleotide identity (ANI), and gene presence–absence analyses revealed three genetically distinct lineages. All strains harbored conserved biosynthetic gene clusters (BGCs) for quorum sensing, carotenoid production, siderophores, and thiopeptides. In contrast, two phosphonate BGCs, key determinants of onion pathogenicity, exhibited lineage-specific distributions. Onion-associated strains from Lineages 1 and 2 carried the Halophos BGC associated with onion tissue necrosis, and onion isolates encoded the *alt* gene cluster conferring thiosulfinate tolerance. Lineage 3 strains, isolated from both onion and rainwater, either lacked a phosphonate BGC loci or carried the HiVir phosphonate BGC. In addition, Lineage 3 strains lacked the *alt* cluster altogether. The localization of these virulence genes in the genome varied, with Halophos integrated in the chromosome, HiVir encoded on the conserved Large *Pantoea* Plasmid, and *alt* located on a small, variable plasmids (plasmid B). The Type IV and Type VI secretion systems showed variable genomic architectures, with plasmid-borne T4SSs and two chromosomal T6SS loci differing in conservation and gene content. Additionally, conserved Pantailocin phage islands were detected in most genomes. Overall, this study reveals that while core metabolic and competitive traits are conserved across *P. allii*, virulence-associated loci display lineage-specific partitioning, reflecting ecological differentiation and evolutionary plasticity within the species.

**Impact Statement:** This study presents a comprehensive comparative genomic analysis of available *Pantoea allii* genomes, a known onion pathogen. By analyzing 38 strains isolated from plant and water sources across three continents, we uncovered lineage-specific distributions of key virulence genes, alongside conserved genetic traits associated with competition and environmental resilience. These findings clarify the genetic basis of *P. allii* pathogenesis and highlight its potential as a biocontrol agent, offering broader insights into how ecologically significant loci contribute to the dual roles of plant-associated bacteria as both pathogens and beneficial microbes.

**Data Summary:** PENDING

## Introduction

The genus *Pantoea* (order Enterobacterales) comprises a group of ubiquitous bacteria capable of colonizing a broad range of ecological niches, including plants, insects, soil, and aquatic environments [1, 2]. Members of this genus are well known for their diverse interactions with plants, which range from mutualistic relationships as plant growth promoting endophytes and epiphytes to antagonistic interactions as plant pathogens [3, 4]. This ecological flexibility enables *Pantoea* species to persist in both natural and agricultural ecosystems, where they contribute to microbial communities and influence plant health.

Among these species, *Pantoea allii* is relatively understudied despite its ecological and agricultural relevance. First described in South Africa from onion (*Allium cepa*) tissues and seeds [5], *P. allii* has been implicated in the Onion Center Rot disease complex alongside *P. ananatis*, *P. agglomerans*, and *P. stewartii* subsp. *indologenes* [6–9]. This disease complex, which causes foliar necrosis and internal bulb decay [6], affects onions during cultivation and post-harvest storage [7] and remains a persistent threat to onion production worldwide. Despite its involvement in disease, *P. allii* is infrequently isolated, and the genomic basis of its pathogenesis and ecological flexibility remain poorly characterized.

Several virulence-associated loci have been described in onion-pathogenic *Pantoea* species. These include the phosphonate biosynthetic clusters HiVir [10–12] and Halophos [13], which encode phytotoxins responsible for inducing onion tissue necrosis, as well as the *alt* gene cluster [14], which confers tolerance to onion-derived antimicrobial thiosulfinates. While these loci are well documented in *P. ananatis* [10], *P. stewartii* subsp. *indologenes* [13], and *P. agglomerans* [12], their distribution and functional conservation in *P. allii* are currently unclear.

Beyond plant pathogenicity, interbacterial competition plays a central role in the ecological success of many bacterial species including *Pantoea* [15]. The Type VI Secretion System (T6SS) is a contact-dependent molecular weapon used to inject antibacterial effectors (toxins) into neighboring cells and has been shown to modulate both microbial competition and host colonization [16, 17]. Genomic analyses of *P. ananatis* and *P. agglomerans* have revealed multiple T6SS gene clusters in each species [18, 19]. In both species, the T6SS-1 cluster encoded a functional system involved in interbacterial antagonism [16, 17], and in the case of *P. ananatis*, it also contributed to plant pathogenicity [17]. In addition to the T6SS, *Pantoea* species deploy tailocins, phage tail-like bacteriocins that mediate targeted and efficient elimination of closely-related competitors [20]. These systems have shown promise as tools for targeted microbial control in agricultural settings. While well-characterized in related species, the distribution and potential ecological roles of T6SS and tailocins in *P. allii* have not been systematically evaluated.

In this study, we performed a comparative genomic analysis of 38 *P. allii* strains isolated primarily from plant hosts and rainwater. *Pantoea* species have previously been reported to exhibit ice nucleation activity [21], a trait that can facilitate survival and dissemination in cold and aquatic environments. Although ice nucleation has not been confirmed in *P. allii*, its recovery from rainwater suggests that similar adaptations could support its persistence and spread. Accordingly, our objectives were to (i) investigate the phylogenomic structure of this infrequently isolated species, (ii) characterize the distribution and diversity of virulence- and competition-associated loci, and (iii) assess phenotypic traits related to onion pathogenicity and ice nucleation. This work advances our understanding of *P. allii* as both a plant-associated pathogen and an environmentally adapted competitor, and it identifies genetic features that may be leveraged for sustainable disease management.

## METHODS

### Culturing

The pure cultures of *Pantoea* alii stored in 10 % glycerol at −80 °C were routinely cultured overnight at 28 °C in Lysogeny broth (LB) or LB agar (10 g/L tryptone, 5 g/L yeast extract, 10 g/L sodium chloride or 15 g/L agar). For ice nucleation assay, strains of *P. allii* were cultured on Reasoner’s 2 Agar (R2A) at 28 °C for 24 hours.

### DNA extraction, whole genome sequencing and genome assembly

20 *Pantoea allii* strains were sequenced for this study (Table 1). The bacterial cells were pelleted by centrifuging 3 ml overnight culture at 2,500 relative centrifugal force (Eppendorf 5810 R) and genomic DNAs were extracted from the cell cultures using commercial DNA extraction kits.

**Table 1.**
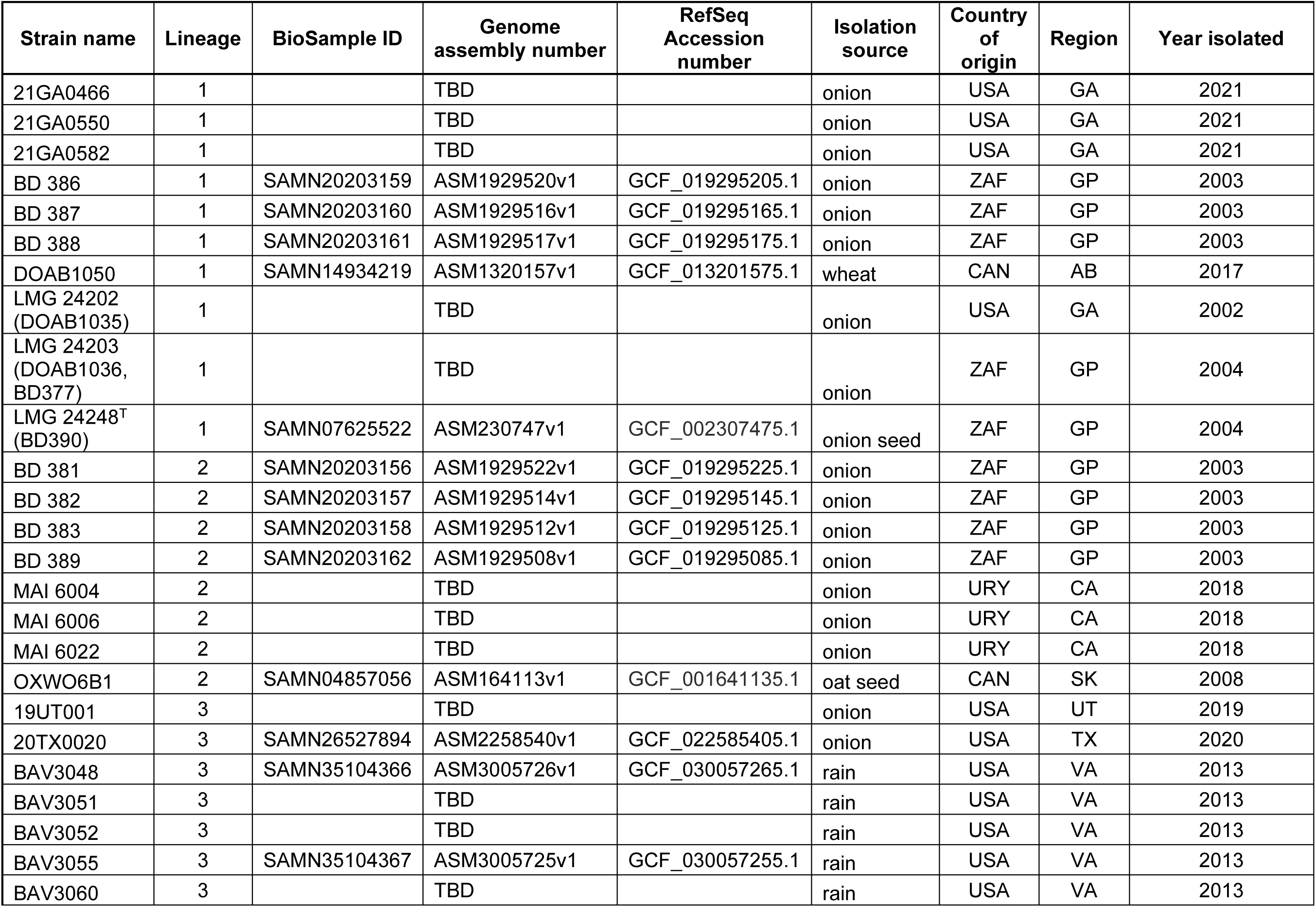
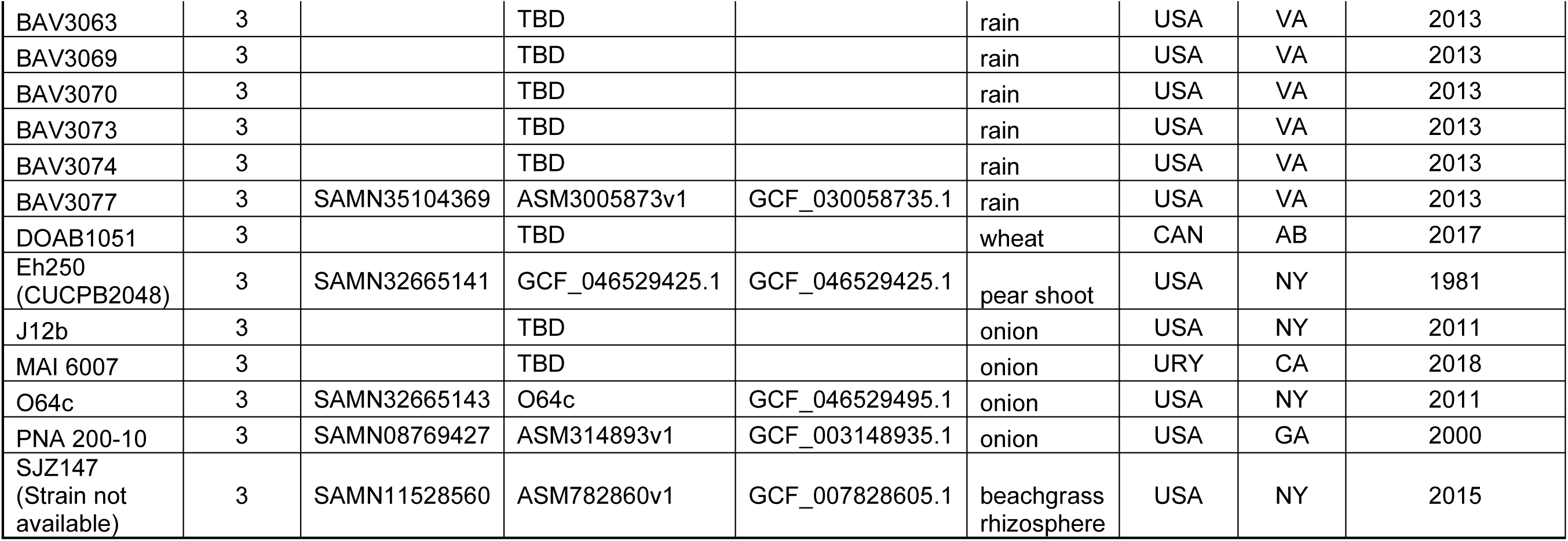
A list of strains used in this study.

Bacterial whole genome library preparation and sequencing were performed by SeqCenter (Pittsburgh, USA) using the Illumina DNA Prep kit with IDT 10bp UDI indices. Sequencing was carried out on an Illumina NextSeq 2000, generating 2×151bp paired-end reads. The raw reads were quality-filtered and adapter-trimmed using Trimmomatic (v3.39) [22]. *De novo* assembly of the processed reads was performed with SPAdes (v3.15.4) [23], and the quality and completeness of the genome assemblies were evaluated using QUAST (v5.0.2) [24] and BUSCO (v5.2.2) [25], respectively.

Genomes were closed by performing hybrid assembly of long and Illumina reads. Long-read sequencing was performed using a MinION Mk1B device (Oxford Nanopore Technologies) equipped with an R9 Spot-ON flow cell. Library preparation was carried out with the Rapid Barcoding Kit (SQK-RBK), and the flow cell was primed using the EXP-FLP002 Flow Cell Priming Kit, following the manufacturer’s protocols. Basecalling of raw signal data was conducted using Guppy (Oxford Nanopore Technologies), and demultiplexing of barcoded reads was performed with qcat (https://github.com/nanoporetech/qcat). Low-quality reads and adapters were removed using FiltLong (https://github.com/rrwick/Filtlong), and high-quality reads were randomly subsampled to 50× coverage, assuming a 5 Mb genome.

Nanopore reads were initially trimmed and filtered for quality using FiltLong (https://github.com/rrwick/Filtlong). A subset of high-quality reads was randomly sampled to a defined coverage depth to standardize input for downstream assembly. Multiple draft assemblies were generated using distinct long-read assemblers, including Flye [26], MiniPolish [27], and Raven [28]. These assemblies were then reconciled into a single consensus genome using Trycycler [29], which integrates structural variation across assemblies to produce a refined consensus. The resulting genome was polished in two stages: first with Medaka (https://github.com/nanoporetech/medaka) using the Nanopore reads, followed by additional polishing with NextPolish [30] using Illumina short reads that had been quality-filtered with FASTP [31]. Final curation steps were performed with custom scripts, which included removal of contaminant sequences (e.g., phiX174), renaming of contigs based on descending sequence length, and reorientation of sequences to place key replication genes, such as *dnaA*, near the beginning of the positive strand.

Genome quality was assessed through annotation with the NCBI Prokaryotic Genome Annotation Pipeline (PGAP) [32] and evaluation of proteome completeness using BUSCO [25]. To validate assembly accuracy, an independent *de novo* assembly was also generated using Unicycler [33] and compared with the consensus assembly using ‘dnadiff’ from the MUMmer package.

### Pangenome analysis and construction of recombination-free core-genome phylogenetic tree

In addition to 20 newly assembled genomes of this study, 18 additional *P. allii* genomes were downloaded from NCBI for pangenome analysis. The strain name, metadata and the GenBank accession numbers of genomes used in the analysis are listed in Table 1. The total of 38 genomes were annotated using PROKKA (v1.14.5) [34] and the resulting annotation files (gff) were analyzed for the pangenome of *P. allii* using Roary (v3.13.0) [35]. The Roary was run with options -e --mafft to generate an alignment of core genes (present in > 95% of genomes) which was used to construct core-genome phylogenetic tree using FastTree v2.1.11 [36]. To build a recombination-free phylogenetic tree, recombination regions of core genes were identified using ClonalframeML (v1.12) [37] and the regions were masked by running cfml-maskrc script (https://github.com/kwongj/cfml-maskrc) prior to reconstructing recombination-free tree.

### ANI, dDDH analysis and LIN assignment

Nucleotide sequences of 38 *P. allii* genome assemblies were analyzed for their genome distance to each other. Pairwise average nucleotide identity (ANI) was calculated using FastANI (v1.33) [38] with default settings and resulting ANI matrix was visualized as a heatmap using a R script available at https://github.com/spencer411/FastANI_heatmap. Digital DNA-DNA hybridization (dDDH) between the type strain *P. allii* LMG24248^T^ (GCF_002307475.1) and 38 genomes of this study was conducted using Genome-to-Genome Distance Calculator v3.0 through Type Strain Genome Server (TYGS) (available at ggdc.dsmz.de/ggdc.php#). The dDDH values calculated from Formula 2 (sum of all identities found in HSPs divided by overall HSP length) were selected. Life Identification Number (LIN) was assigned to each genome through LINbase webserver (http://www.LINbase.org).

### Identification of secondary metabolite biosynthetic gene clusters and ice nucleation gene

Secondary metabolite biosynthetic gene clusters (BGCs) in the genomes of *P. allii* were determined using antiSMASH included in the Beav (v1.4.0) pipeline [39]. Previously characterized HiVir and Halophos biosynthetic gene clusters (BGCs) responsible for producing onion necrosis factors in *P. ananatis* [10] and *P. stewartii* subsp. *indologenes* [13] and ice nucleation protein encoding gene of *P. ananatis* PNA 97-1 [40] were searched against 38 *P. allii* genomes using nucleotide BLAST+ (v2.14.1) search [41]. Presence and absence data of virulence gene clusters were included in the metadata table (Table S1).

### Identification of type secretion systems

The presence of type III, type IV and type VI secretion systems were identified using Beav v1.4.0 pipeline [39]. A type secretion system was deemed present when a full quorum of core or mandatory genes of a system was met. For draft genomes, a quorum of mandatory genes was relaxed by one. Conjugative elements were further screened from Prokka protein annotations of 38 *P. allii* genomes by MacSyFinder [42] using CONJScan models [43].

### Detection of Phage and Pantailocin Islands

Genome-wide detection of phage genes and islands was performed on 38 *Pantoea allii* genomes using PHASTEST (https://phastest.ca/). The Pantailocin island, previously characterized in *P. ananatis* ATCC 35400 and *P. stewartii* subsp. *indologenes* ICMP 10132 [20], is flanked by the *rpoD* and *sulP* genes. This region was extracted from the nine closed *P. allii* and compared to those of *P. ananatis* and *P. stewartii* by clinker analysis described below.

### Gene cluster homology and synteny analysis

Type VI secretion system loci and Pantailocin gene clusters from nice closed-genome *Pantoea allii* strains were extracted in GenBank format for comparative analysis. Type VI loci were compared to those of *P. ananatis* DZ-12 (GCA_003849975.1) and *P. agglomerans* pv. *betae* strain 4188 (GCF_001662025.2), while Pantailocin gene clusters were compared to those of *P. ananatis* ATCC 35400 (GCF_029433965.1) and *P. stewartii* subsp. *indologenes* ICMP 10132 (GCF_029433915.1), using Clinker (https://cagecat.bioinformatics.nl/tools/clinker) with default settings [44]. Collinearity and homology of plasmid Bs from closed genome *P. allii* strains were compared using MAUVE alignment tool [45].

### Replicon comparison by BRIG

The genome similarity of nine closed genome strains was compared by replicons using BRIG (v0.95) [46]. *Pantoea allii* LMG24248^T^ was used as a reference sequence (central ring) for chromosomes whereas Eh250 was used as a central ring for Large *Pantoea* Plasmid (pLPP) and plasmid B. The genomic location of HiVir, Halophos and *alt* (thiosulfinate tolerance) gene clusters were manually searched and indicated on the ring diagrams.

### Red onion scale necrosis assay

Single bacterial colonies were inoculated into 3 ml of LB broth and incubated overnight at 28 °C with shaking at 200 rpm. To prepare the bacterial inoculum for the red onion scale necrosis (RSN) assay, the overnight cultures were pelleted and resuspended in sterile 1X phosphate-buffered saline (PBS) to an optical density (OD) of 0.3 at 600 nm. The RSN assay was performed according to the previously described protocol [47]. The onion scales were visually assessed and imaged at 4 days post-inoculation (dpi). Two independent RSN assays were conducted.

### Ice nucleation assay

Ice nucleation activity was assessed using a droplet freeze assay adapted from Vali, 1971 [48]. A bacterial suspension with an OD_600_ of 0.2 (approximately 10^8^ CFU/mL) was prepared in molecular grade water. The suspensions were incubated at 4 °C for at least 1 hour. Thirty 20 µL droplets were then placed onto a parafilm boat floating on a glycerol bath maintained at −2 °C in a cooling thermostat (Lauda Alpha RA24, LAUDA-Brinkmann, Delran, NJ, USA). The droplets were cooled from −2 °C to −12 °C at a rate of −1 °C every ten minutes. The temperature at which freezing onset occurred and the temperature at which 50 % of the droplets froze (T_50_) were recorded.

## RESULTS

### *Pantoea allii* forms three genetically distinct lineages with high genomic similarity

Genetic diversity and phylogenetic relationships among environmental and plant-associated *P. allii* strains were investigated by analyzing 18 publicly available and 20 newly sequenced genomes. BUSCO analysis indicated 100% completeness of the newly sequenced genomes (Table S2). The pangenome of the 38 *P.* allii strains comprised of a total of 10,127 orthologous and 2,950 unique genes. Among these 3,700 were core genes (present in 99 to 100% of genomes), 87 were soft-core genes (95 to 99%), 1,400 were shell genes (15 to 95%) and 4,940 were cloud genes (0 to 15%). The average number of core genes per *P. allii* genome in this study was 3,787 while the total gene count averaged 4,591. The 38 *P. allii* strains shared average nucleotide identity (ANI) ranging from 98.33% to 100% (Figure S1) and digital DNA-DNA hybridization (dDDH) values ranging from 89.1% to 100% (Table S3). The genome size varied from 4.7 Mb to 5.2 Mb with an average GC content of approximately 53 % (Table S4).

A phylogeny based on the core-genome (n= 3,700) revealed three distinct *P. allii* lineages which were consistent with the LINgroups assigned by LINbase (Lineage 1 = 51_A_6_B_1_C_0_D_1_E_0_F_0_G_0_H_0_I_0_J_0_K_; Lineage 2 = 51_A_6_B_1_C_0_D_1_E_0_F_0_G_0_H_0_I_0_J_1_K_; Lineage 3 = 51_A_6_B_1_C_0_D_1_E_0_F_0_G_0_H_0_I_0_J_2_K_) (Figure 1, Table S5). These lineages were also supported by the groupings based on ANI (Fig. S1) and gene presence-absence profiles (Fig. S2). Strains belonging to Lineages 1 and 2 were isolated from various countries including South Africa, Uruguay and the state of Georgia whereas Lineage 3 was comprised strains predominantly isolated from within the United States.

**Figure 1.**
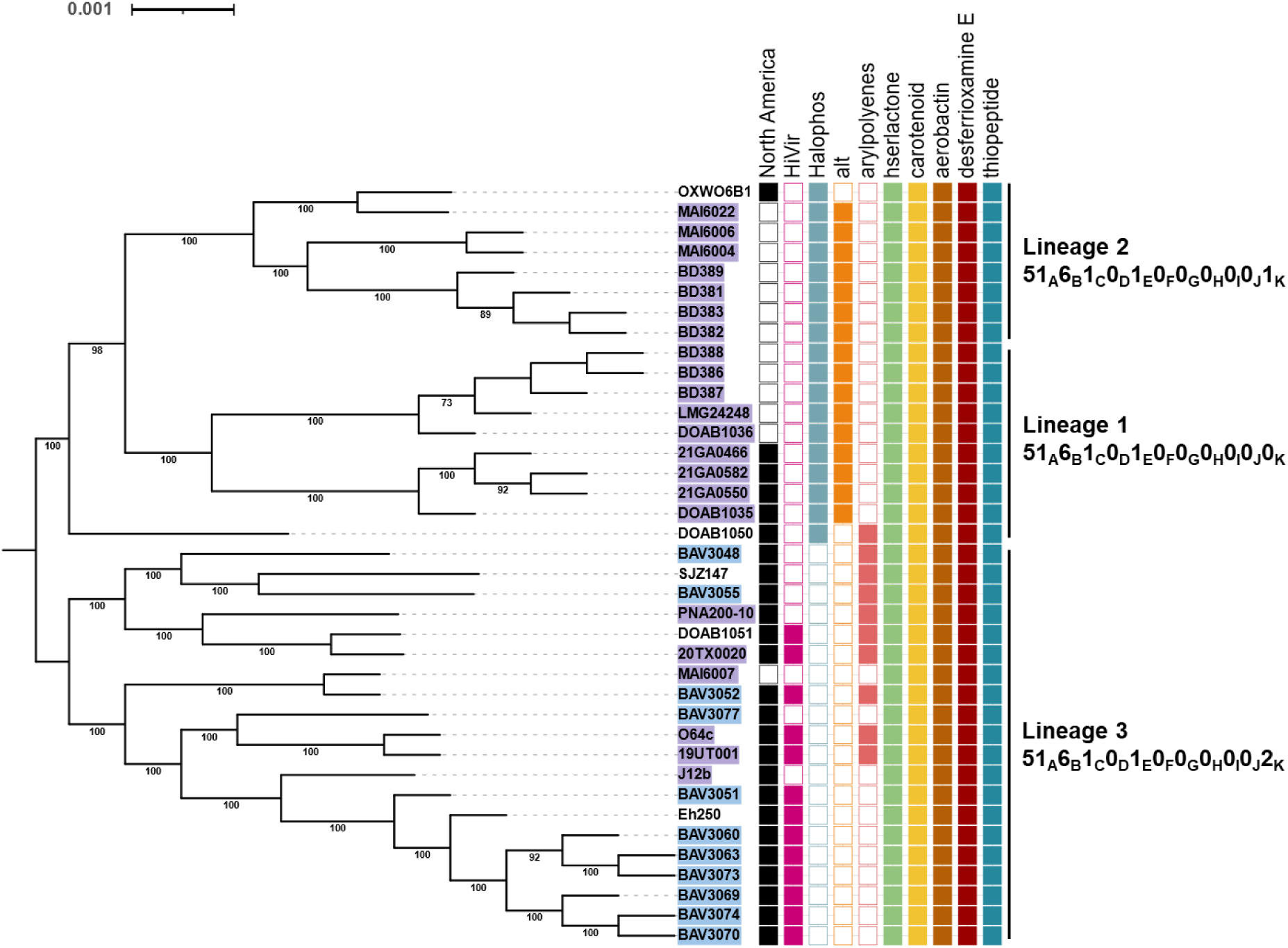
Three distinct lineages are present in *Pantoea allii*. A recombination-free, approximately maximum-likelihood phylogenetic tree constructed based on 3,700 core genes shared among 38 *Pantoea allii* strains using FastTree (v2.1.11) and ClonalFrameML (v1.12). The general time reversible (GTR) nucleotide substitution model was selected as the best-fit evolutionary model and the robustness of groupings was assessed by the approximate likelihood-ratio test (aLRT). The tree was midpoint-rooted and edited in iTOL (v6.8, available at https://itol.embl.de/). Strain names are highlighted in blue (rainwater) and purple (onion-associated) to indicate their isolation source. The colored boxes next to strain names indicated the geographic origin (North America) and the presence of virulence genes (HiVir, Halophos), antimicrobial resistance genes (*alt*, *<u>al</u>*licin *t*olerance), and biosynthetic gene clusters (BGCs) for pigment (aryl polyenes, carotenoid), quorum sensing (acylated homoserine lactone), siderophores (aerobactin, deferrioxamine E) and antimicrobials (thiopeptide). LINgroup numbers are shown for each lineage.

### Presence of broadly conserved secondary biosynthetic gene clusters

Analysis of 38 *P. allii* genomes using antiSMASH [39] identified secondary metabolite biosynthetic gene clusters (BGCs) with more than 70% sequence homology to known homoserine lactone, aryl polyene, carotenoid, aerobactin, desferrioxamine E, and thiopeptide clusters (Table S6). With the exception of aryl polyene, all identified BGCs were conserved across the three *P. allii* lineages (Fig. 1). The homoserine lactone BGC in *P. allii* contained the quorum sensing genes *esaI* acyl-homoserine lactones (AHL) synthase and *esaR* transcription factor which mediate the synthesis of and regulation by AHLs quorum sensing molecules [49]. Carotenoids BGCs which encode lipid-soluble pigments involved in photoprotection and oxidative stress resistance, were universally identified on the Large *Pantoea* Plasmid [50, 51]. These clusters included a conserved *crt* operon (*crtE*, *crtX*, *crtY*, *crtI*, *crtB* and *crtZ*). In contrast, aryl polyene BGCs, which encode structurally similar pigmented metabolites but are synthesized via type II polyketide synthases [52] were only detected in a subset of *P. allii* strains belonging to Lineage 3. Additionally, two siderophore BGCs resembling those of aerobactin and desferrioxamine E were identified [53], suggesting presence of iron scavenging mechanism in *P. allii*. A BGC encoding a ribosomally synthesized and post-translationally modified thiopeptide was also detected [54], indicating the potential for antimicrobial compound production in *P. allii*.

### Lineage-specific distribution of phosphonate biosynthetic and thiosulfinate tolerance (*alt*) gene clusters

To date, two phosphonate BGCs have been functionally characterized for their role in the pathogenicity of *Pantoea* species in onion. These BGCs, named Halophos [13] and HiVir [10], encode phytotoxic phosphonate compound that induces necrosis in onion tissue. Halophos was exclusively identified in *P. allii* strains from Lineage 1 and 2, which are predominantly onion-associated, with the exceptions of strains DOAB1050 and OXWO6B1, isolated from wheat and oat seed, respectively (Table 1). The Halophos-positive strains were collected from North and South America and Africa. In contrast, HiVir was found in Lineage 3 strains, isolated from rainwater, onion, pear shoot and wheat in the United States. Notably, six strains from Lineage 3 (BAV3048, BAV3055, BAV3077, MAI6007, PNA200-10, J12b and SJZ147) lacked both phosphonate BGCs, and no strain was found to contain both gene Halophos and HiVir simultaneously. However, strain MAI6022 harbored an additional phosphonate BGC distinct from HiVir, in addition to Halophos.

The *alt* tolerance gene cluster which encodes enzymes that alleviate thiosulfinate associated thiol stress resulting from onion necrosis [14] was present only in onion-associated, Halophos-positive strains from Lineage 1 and 2. This gene cluster was absent in DOAB1050, a strain isolated from wheat, and OXWO6B1, a potato late blight biocontrol strain which inhibits the growth of *Phytophthora infestans* [55].

### Onion pathogenicity assays reveal phosphonate biosynthetic gene cluster (BGC)-linked necrosis phenotype

To assess the virulence of *P. allii* strains, the pathogenicity of 37 strains was tested using detached fleshy red onion scales, one strain was not available (Fig. 2). Four days post inoculation (dpi), distinct necrosis patterns were observed HiVir-positive *P. allii* strains caused a characteristic oval-shaped white clearing zone on the onion scales, while some Halophos-positive strains, such as BD388, MAI6004 and MAI6022 produced a sunken yellow clearing zone (Fig. 2). In addition to these distinct necrotic patterns, faint discoloration was observed around the inoculation point for certain Halophos-positive strains as well as for strains that lacked both HiVir and Halophos BGCs.

**Figure 2.**
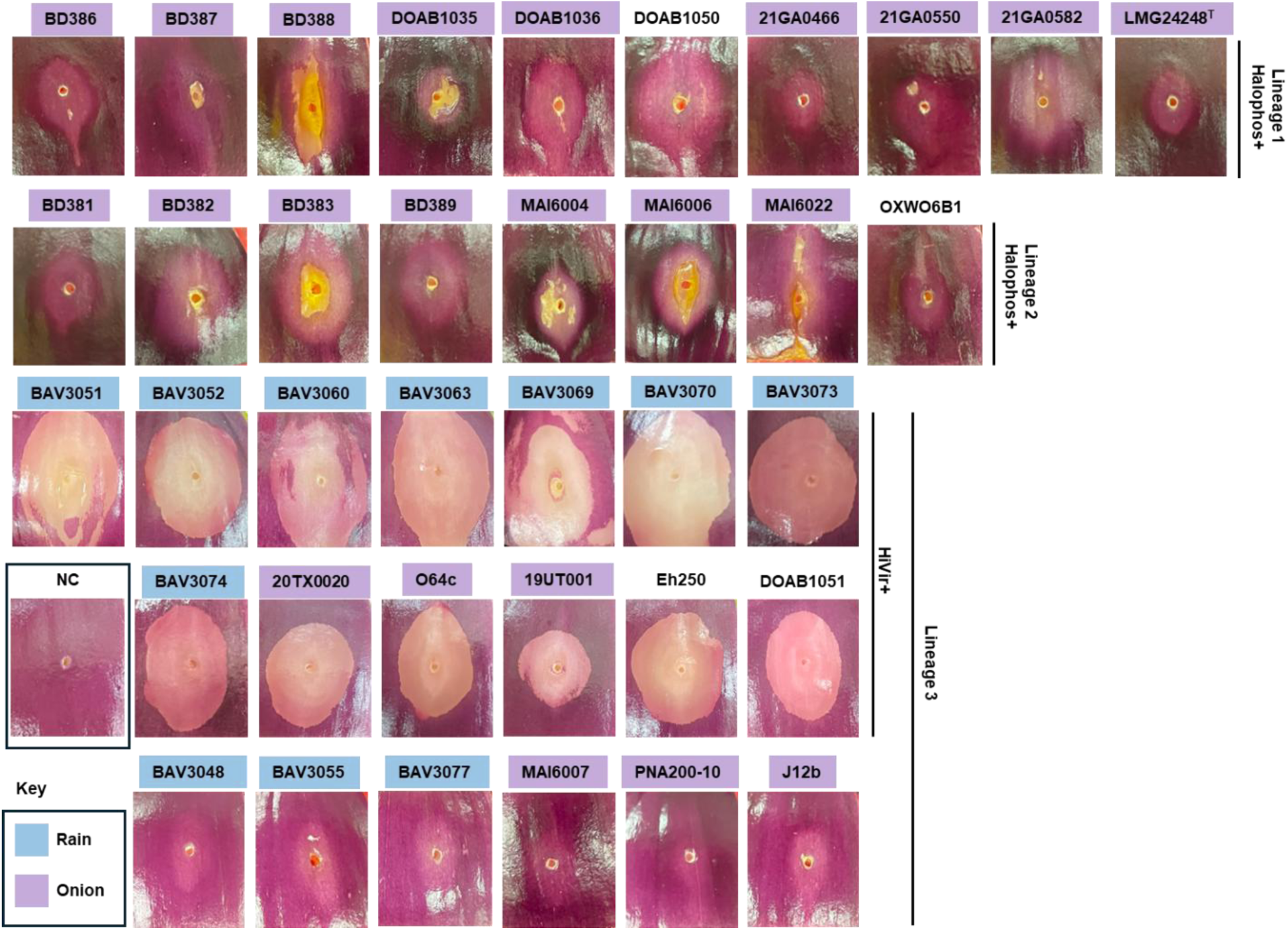
Presence of HiVir and Halophos phosphonate biosynthetic gene clusters (BGCs) is associated with distinct red onion scale necrosis phenotypes. Pathogenicity of 37 *Pantoea allii* strains was assessed using detached red onion scales. Four days post-inoculation (dpi), HiVir-positive strains, isolated from rainwater and onion tissues and belonging to Lineage 3, caused white clearing (necrotic) zones. In contrast, some Halophos-positive strains, exclusively from Lineages 1 and 2, produced a yellow, sunken necrosis phenotype. Strains lacking both HiVir and Halophos BGCs did not induce necrosis, although faint lightening of the red onion scales was occasionally observed. No symptoms were observed on scales inoculated with the negative control (NC). Strain names are colored blue (rainwater) and purple (onion-associated) to indicate their source of isolation.

### Halophos, HiVir and *alt*-specific genomic localization on chromosomes and plasmids

To investigate the genomic organization of key onion pathogenicity gene clusters, the location of Halophos, HiVir and *alt* were compared across nine *P. allii* strains with closed genomes. The Halophos was located on the chromosome (Fig. 3A) whereas HiVir was found on the Large *Pantoea* Plasmid which was identified by the presence of carotenoid biosynthesis genes (Fig. 3B). The *alt* gene cluster, associated with thiosulfinate stress tolerance, was present on plasmid B in Halophos-positive strains (DOAB1035, LMG24248 and MAI6022) but absent from plasmid B in HiVir-positive strains (Eh250, MAI6007 and O64c). Plasmid B varied in size, ranging from 113.1 to 174.5 kb, and exhibited substantial differences in gene content between Halophos-positive and HiVir-positive strains (Fig. 3C). Most closed-genome strains (eight out of nine) contained three replicons: the chromosome, the Large *Pantoea* Plasmid, and plasmid B. However, strain MAI6007 harbored an additional replicon, plasmid C (108.3 kb), which contained a distinct set of genes but shared plasmid replication genes and remnants of plasmid B-associated sequences (Fig. 3C).

**Figure 3.**
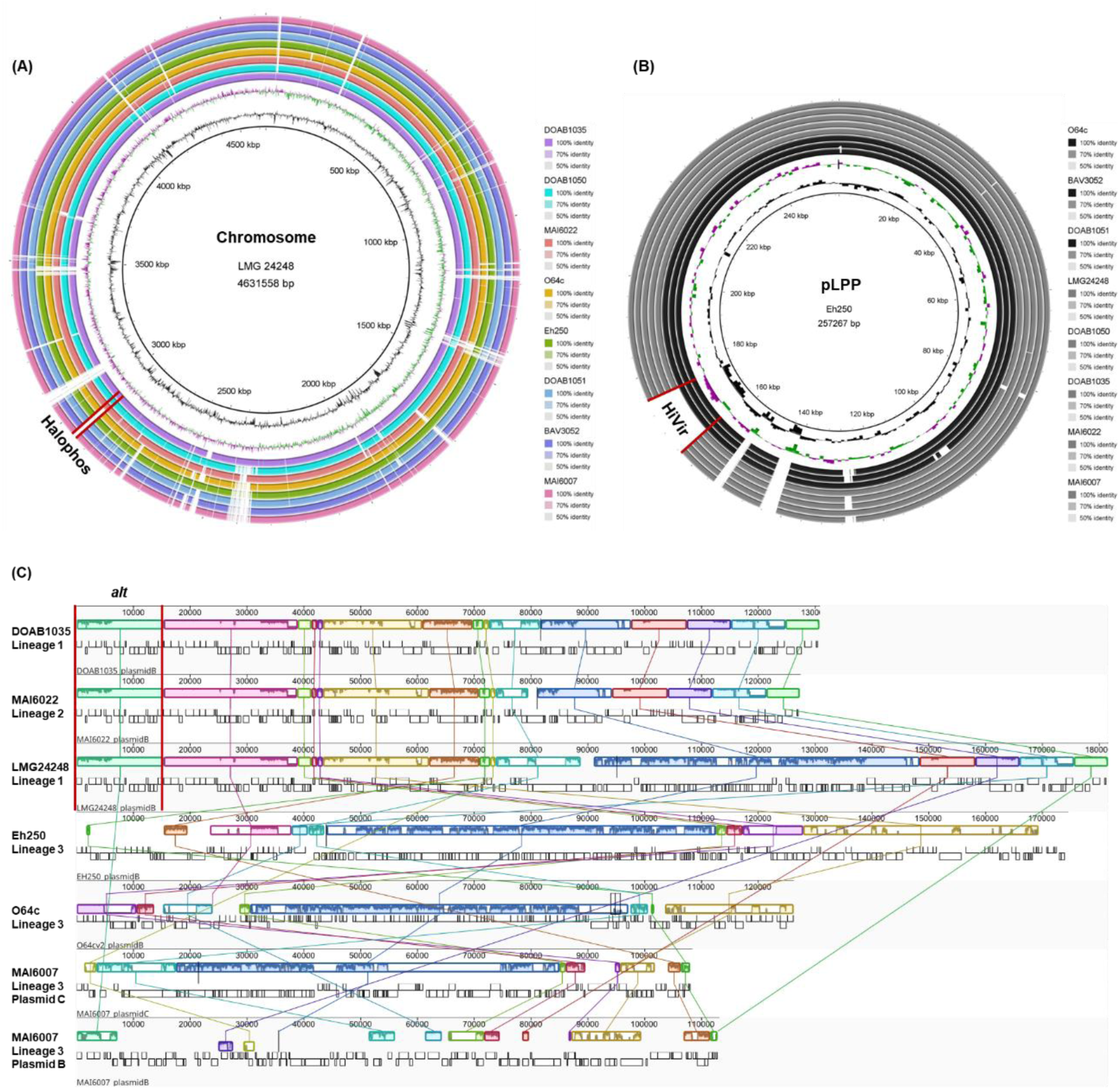
Genomic localization of Halophos, HiVir, and *alt* gene clusters on chromosomes and plasmids. The genomic locations of onion virulence gene clusters (HiVir, Halophos) and the antimicrobial resistance gene cluster (*alt*) were investigated in nine closed-genome *Pantoea allii* strains representing each lineage, using BRIG [46] and MAUVE [45] analyses. The analyses revealed that **(A)** Halophos is located on the chromosome, **(B)** HiVir on the large *Pantoea* plasmid (pLPP), and **(C)** the *<u>al</u>*licin *t*olerance cluster (*alt*) is carried on plasmid B. A high degree of conservation was observed for the chromosome and pLPP1 across strains, while plasmid B was more variable. In Lineages 1 and 2, plasmid B carried the *alt* cluster (mint block) and showed similar sequence content, whereas plasmid B in Lineage 3 strains lacked *alt* and was more conserved within the lineage but divergent from those of Lineages 1 and 2. Strain MAI6007 carried an extra replicon, plasmid C shared conjugate type IV secretion system genes (dark blue block) on plasmid Bs of LMG24248^T^, Eh250 and O64c.

### Presence of Type IV and Type VI Secretion Systems in *P. allii*

Secretion systems often play a crucial role in bacterial interactions with hosts and other microbes, influencing virulence, competition, and environmental adaptation. To assess the presence and distribution of secretion systems in *P. allii*, the 38 genomes were analyzed using Beav (v1.4.0) [39]. While the Hrp-type Type 3 Secretion System (T3SS), typically associated with plant pathogenicity, was absent, genes encoding conjugative Type 4 Secretion System (T4SS) and Type 6 Secretion System (T6SS) were identified on plasmids and chromosomes of *P. allii*, respectively (Table S7). To further investigate the gene synteny and architecture of T4SS and T6SS clusters, nine closed genome strains were selected for MAUVE [45] and clinker [44] analyses, respectively. Conjugative T4SS genes were primarily localized on plasmid B, represented by the dark blue block in the MAUVE alignment (Fig. 3C). Beav’s TXSS scan (Table S7) and CONJScan by MacSyFinder [42] predictions indicated that plasmid B of strains LMG24248, O64c, Eh250 carried a structurally complete T4SS cluster, whereas plasmid C of MAI6007 also harbored a complete T4SS (Table S8). In contrast, no T4SS genes were detected on plasmid B of DOAB1035 and MAI6022, nor on the Large *Pantoea* Plasmid (LPP) of any *P. allii* strain.

Type 6 Secretion Systems (T6SS) have been found to play a key role in bacterial competition and interactions with plant hosts. In *P. allii*, two chromosomal T6SS loci, T6SS-I and T6SS-II, were identified across the analyzed genomes. T6SS-I was positioned downstream of an EmmdR/YeeO family multidrug/toxin efflux MATE protein (LMG24248 locus tag: PENDING) while T6SS-II was located downstream of a glycerophosphodiester phosphodiesterase family protein (LMG24248 locus tag:PENDING). These T6SS loci were flanked by tRNA-Asn and tRNA-Arg, respectively. Analysis using Beav’s TXSS scan predicted that T6SS-I contained structurally complete core gene cluster (Table S7, Fig.4A), while T6SS-II exhibited varying degrees of completeness across strains (Table S7, Fig. 4B). Among the nine closed genome strains, only MAI6007 and BAV3052 harbored a complete T6SS-II gene cluster (∼52 kb). In contrast, O64c (10.1 kb) and DOAB1050 (14.1 kb) contained the shortest T6SS-II loci, while DOAB1051 (76.5 kb) and DOAB1035 (77.8 kb) had the largest. The expansion of T6SS-II in DOAB1051 was attributed to the integration of phage-related genes, whereas in DOAB1035, the presence of genes encoding hypothetical proteins, integrases, transposases, and plasmid replication proteins contributed to its increased size, (Fig. 4B). Notably, plasmid replication genes were also detected within the T6SS-II loci of MAI6022 and DOAB1035 (Fig. 4B).

**Figure 4.**
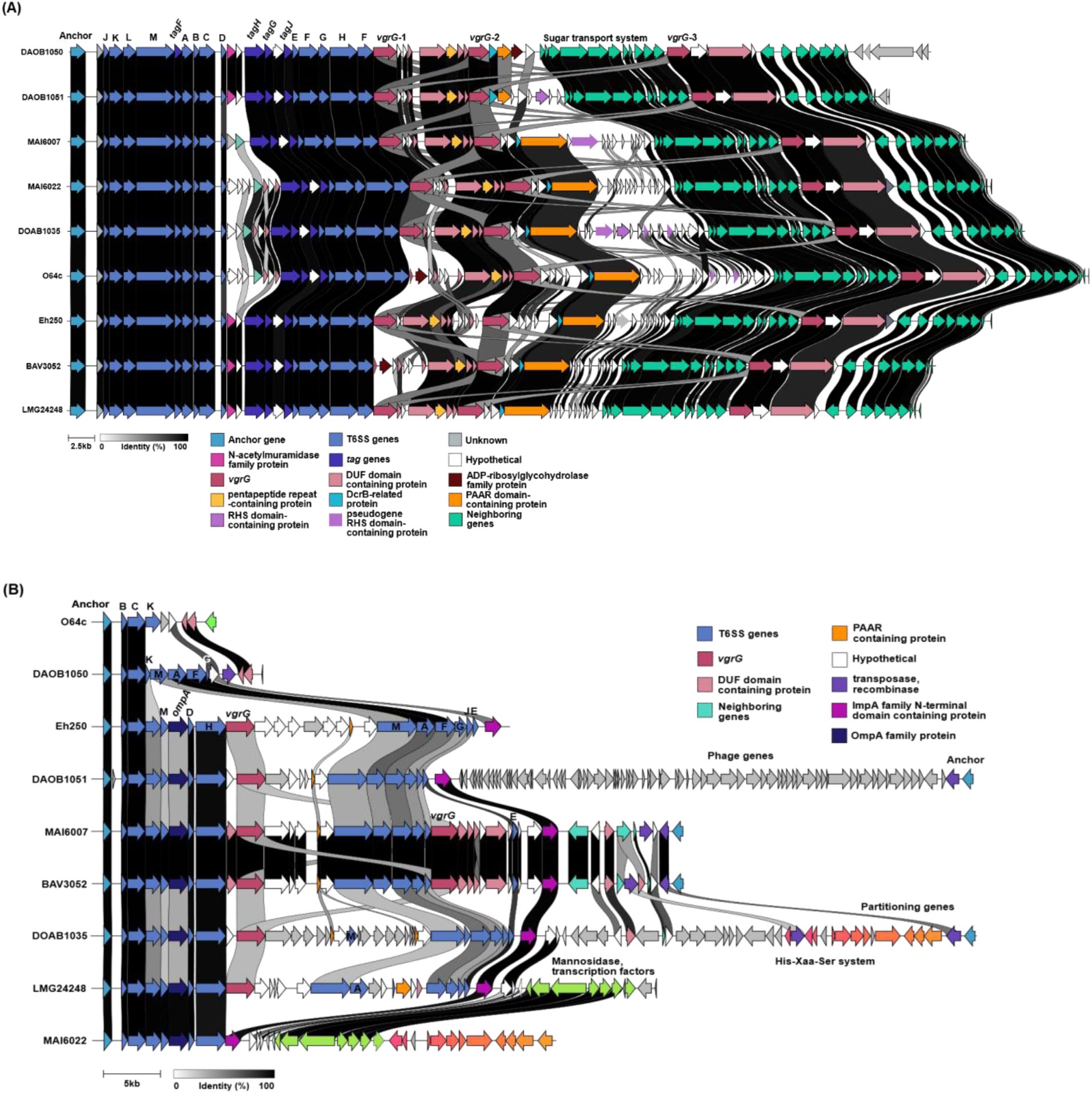
Comparative analysis of two chromosomal T6SS loci in *Pantoea allii* reveals differences in gene content conservation. Two chromosomal Type VI Secretion System (T6SS) loci were identified in nine closed genome *P. allii* strains and compared using clinker [44] to assess gene content and synteny. Genes are represented as arrows and colored by predicted function. **(A)** T6SS-I included a complete set of conserved core structural genes [*tssA–M*, *hcp* (*tssD*), *vgrG* (*tssI*), and *clpV* (*tssH*)] arranged in a consistent synteny across all strains, while **(B)** T6SS-II was high variable in structural gene content among strains. Only MAI6007 and BAV3052 possessed a nearly complete set of core T6SS genes and other strains contained partial clusters with a reduced number of core structural genes.

Compared to T6SS-II, T6SS-I exhibited greater conservation across *P. allii* strains. Downstream of the structural and T6SS-associated (tag) genes, three distinct islands consist of *vgrG* (spike protein encoding gene), and *vgrG*-associated genes were identified, encoding hypothetical and domain of unknown function (DUF)-containing proteins (Fig. 4A, Table S9). However, in strains O64c and BAV3052 (Figure 4A), the first *vgrG* gene was predicted to be a pseudogene. Despite this, the downstream genes encoding pentapeptide repeat (COG1357), DUF3540 (pfam12059) and DUF4150 (pfam13665) domain-containing proteins remained intact in all nine closed genomes. The second *vgrG* (*vgrG*-2) island contained the highest number of hypothetical protein-encoding genes and was enriched in genes that encode RHS domain- or repeat-containing proteins (Fig. 4A, Table S9). In DOAB1035 and MAI6007, some RHS domain-containing protein encoding genes were predicted to be pseudogenes (Fig. 4A, Table S9). Among the conserved genes on this island, DcrB-related and PAAR (also DUF6531) domain-containing protein encoding genes were consistently present. The PAAR domain-containing proteins also carried an RHS domain or element. In DOAB1050, PAAR domain-containing protein also harbored a domain (pfam03496) found in actin-ADP-ribosylation toxin followed by an ADP-ribosylglycohydrolase family protein. The third *vgrG* island (*vgrG*-3) was located downstream of genes encoding PTS fructose transporter and ABC transporter proteins and were most highly conserved *vgrG* island of the three, consisting of DUF6531domain-containing and hypothetical protein encoding genes (Fig. 4A, Table S9).

### Presence of Pantailocin islands in *P. allii*

Prophage islands were identified in the genomes of 38 *P. allii* strains using PHASTEST [56], which assessed their completeness based on phage structural genes (Table S11). Most strains contained at least one intact prophage island. However, no ‘intact’ prophage islands were detected in the draft genomes of BAV3051, 21GA0446, 21GA0550, 21GA0582, and MAI6006. A manual alignment and clinker analyses of intact prophage islands in nine closed genomes revealed a conserved genomic context. These islands ranged in size between 35 kb to 71 kb were consistently flanked by *rpoD* (locus tag: PENDING) and *sulP* (locus tag: PENDING) genes (Fig. 5). Additionally, they shared homology with the Pantailocin island previously identified in *P. ananatis* ATCC 35400 and *P. stewartii* subsp. *indologenes* ICMP10132 [20]. Interestingly, in DOAB1050 and Eh250, the Pantailocin island closely resembled that of *P. stewartii* subsp. *indologenes* ICMP10132, an expanded phage island compared to its counterpart in *P. ananatis* ATCC 35400. In contrast, the Pantailocin islands in BAV3052, DOAB1035, DOAB1051, LMG24248, MAI6007, MAI6022, and O64c were more similar to that of *P. ananatis* ATCC 35400 which exhibited partial homology to the Pantailocin islands in DOAB1050, Eh250 and *P. stewartii* subsp. *indologenes* ICMP10132. Additionally, DOAB1051 Pantailocin island contained extra phage genes (gray arrows) that lacked homology to rest of phage genes found in *P. allii*, *P. ananatis* ATCC 35400 and *P. stewartii* subsp. *indologenes* ICMP10132 strains, (Fig. 5).

**Figure 5.**
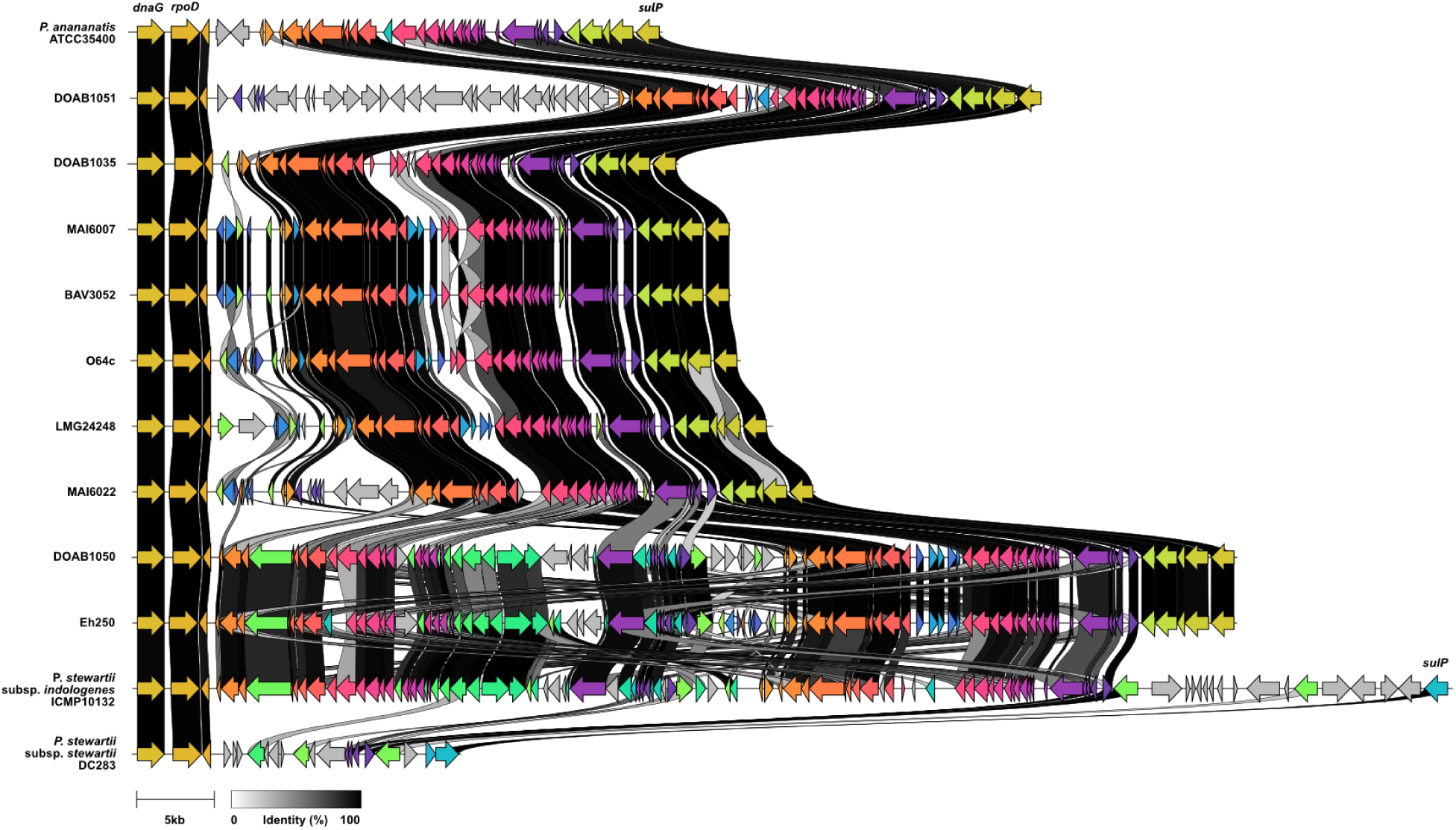
*Pantoea allii* strains harbor the Pantailocin island. The Pantailocin island was initially detected as a prophage island by PHASTEST in *P. allii* strains, located in a conserved chromosomal region flanked by *rpoD* and *sulP* genes. Clinker analysis of the island to the functionally characterized Pantailocin island in *P. ananatis* ATCC 35400 and *P. stewartii* subsp. *indologenes* ICMP10132 revealed high homology. The islands in *P. allii* strains DOAB1035, DOAB1051, BAV3052, O64c, LMG24248, MAI6007, and MAI6022 were most similar to the Pantailocin island found in P. *ananatis*, while those in DOAB1050 and Eh250 were more closely aligned with the island found in *P. stewartii*.

### Widespread ice nucleation activity among *P. allii* strains

Ice nucleation protein encoding gene was detected in all 38 *P. allii* genomes while ice nucleation activity was observed in 24/26 *P. allii* strains tested in this study. 80% of the strains exhibited a freezing onset temperature of −10°C or higher (Table 2). Additionally, over 80% of strains failed to reach T_50_ before the conclusion of the assay. Notably, strain Eh250 displayed the highest activity with a freezing onset temperature of −4.0°C and a T_50_ of −7°C. Strains BD389 and 20TX0020 did not exhibit INA within the temperature range tested, suggesting a potential absence of ice nucleation activity under these conditions. It is possible that this strain exhibits INA at lower temperatures or under different growth conditions [21].

**Table 2.**
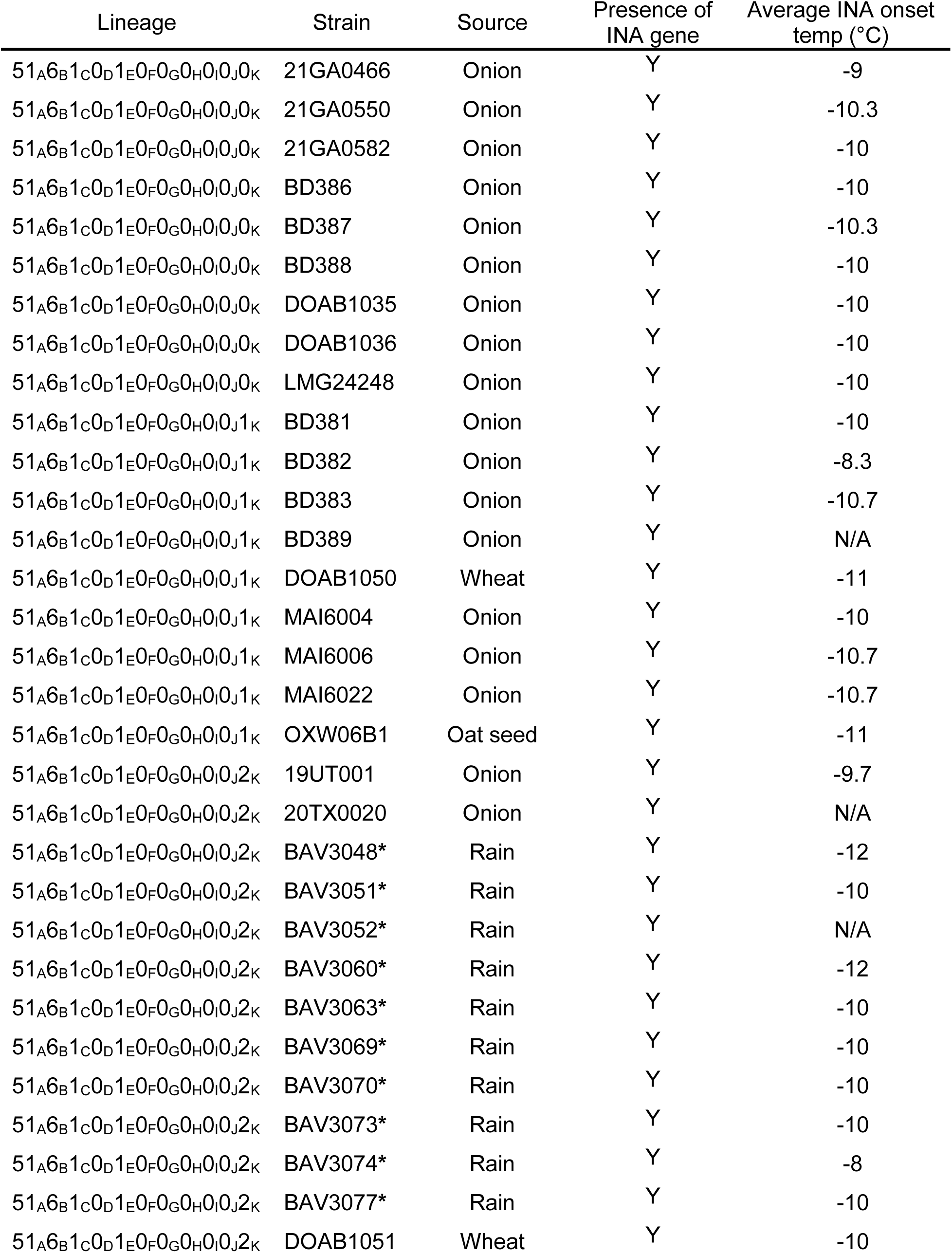
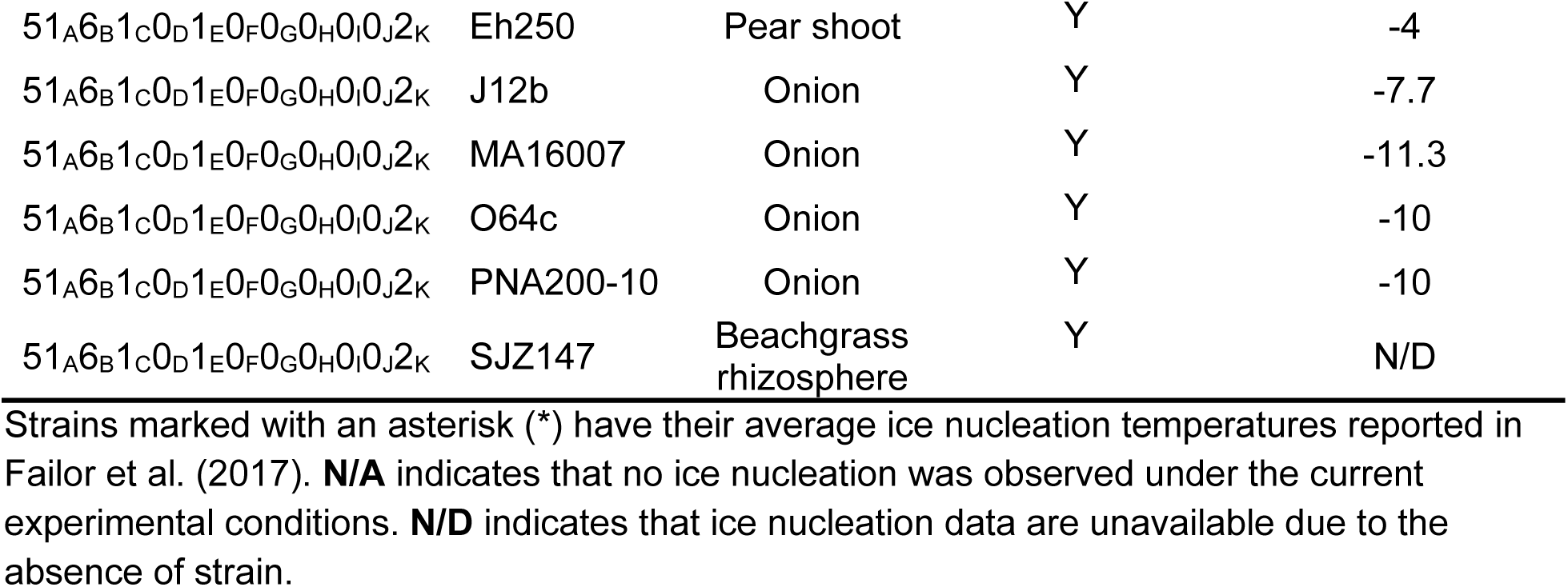
Ice nucleation activity of *Pantoea allii* strains.

## Discussion

*Pantoea* species are broadly distributed in diverse environments and are frequently associated with plants as both epiphytes and endophytes [1, 3, 4]. Several species, including *P. agglomerans*, *P. ananatis*, *P. stewartii* subsp. *indologenes* and *P. allii* are causal agents of Onion Center Rot. While *P. allii* was originally from onion plant and seed [5] and implicated in postharvest bulb rot [7], it is far less frequently isolated from symptomatic onions. Interestingly, a substantial portion of the *P. allii* strains characterized in this study were recovered from rainwater, raising the possibility that this species may be underrepresented in surveys focused exclusively on plant tissue. This observation suggests that *P. allii* may persist primarily in non-plant reservoirs or occupy different ecological niches than other Onion Center Rot pathogens, and that current isolation practices may be missing its primary environmental context.

Despite differences in prevalence and isolation sources, onion-pathogenic *Pantoea* species share common virulence strategies centered on the production of phosphonate phytotoxins that induce necrosis in onion tissue, namely HiVir and Halophos. While HiVir was initially identified on the chromosome of *P. ananatis* [10], it has since been found on the plasmids in *P. agglomerans* [12] and *P. vagans* [57]. In contrast, Halophos has not been reported in these species [12, 58, 59]. However, both HiVir and Halophos have both been identified in strains of *P. stewartii* subsp. *indologenes* [13], and findings from this study indicate that the same is true for *P. allii*. The strains of *P. allii* carrying either the HiVir or Halophos induced distinct necrosis symptoms on detached fleshy red onion scales, however, the regulatory mechanisms governing these gene clusters and the precise molecular targets of their phytotoxins remain unknown.

Phylogenomic analyses grouped geographically diverse *P. allii* strains into three distinct lineages. Strains in Lineages 1 and 2 were distributed across multiple regions, while Lineage 3 was composed primarily of strains from the United States. Interestingly, this phylogenetic structure corresponded with the distribution of key virulence factors. Halophos-positive strains were found only in Lineages 1 and 2, whereas Lineage 3 strains uniformly lacked Halophos but instead carried the HiVir gene cluster. The thiosulfinate tolerance gene cluster (*alt*) was detected exclusively in Halophos-positive strains. In contrast, HiVir-positive strains from Lineage 3 lacked *alt*, despite harboring plasmid B, the same plasmid that carries *alt* in Halophos-positive strains (Fig. 1 and 3). The factors driving this unexpected overlap between phylogeography and virulence factor distribution remain unclear. However, the absence of the *alt* cluster in HiVir-positive *P. ananatis* [60] and *P. agglomerans* [12] has also been observed. In those species, necrosis-inducing HiVir-positive strains may rely on co-occurring *alt*-carrying strains for *in planta* thiosulfinate detoxification. This phylogeographic signal, coupled with the partitioning of virulence traits, raises intriguing questions about the evolutionary forces shaping the distribution, ecology, and host interactions of *P. allii*.

Two genomic loci encoding Type 6 Secretion System (T6SS) gene clusters were identified in *P. allii* strains. In comparison, up to three T6SS gene clusters have been reported in *P. ananatis* [17–19] and two in *P. agglomerans* [16]. In both species, T6SS-1 encodes a complete and functional antibacterial system that has been experimentally characterized. Based on homology, *P. allii* T6SS-I and T6SS-II correspond to T6SS-1 and T6SS-2, respectively, in *P. ananatis* and *P. agglomerans* [16, 17, 19]. Unlike the structurally incomplete T6SS-2 in *P. ananatis* and *P. agglomerans*, *P. allii* strains MAI6007 and BAV3052 harbored computationally predicted complete T6SS-2 (Pall T6SS-II), suggesting potential functional differences. It is also possible that components of the two T6SS clusters in *P. allii* could interact, forming derivatives of the T6SS apparatus. Additionally, the variability of T6SS-I, the presence of plasmid partitioning genes in T6SS-I of DOAB1036 and MAI6022, and the presence of unique phage genes in DOAB1051 (Fig. 4B) suggest that T6SS-I is more prone to recombination and integration events than T6SS-II.

The Type 6 Secretion System (T6SS) gene clusters of *P. allii* are enriched with RHS (rearrangement <u>h</u>ot <u>s</u>pot) domain-containing proteins, which are often associated with polymorphic C-terminal toxin domains [61, 62]. The *vgrG*-1 island of *P. allii* T6SS-I (Fig. 4A) harbors a pentapeptide repeat domain-containing protein (COG1357), a feature also observed in *P. ananatis* DZ-12 (Zhao et al., 2024). In *P. ananatis*, this protein, identified as the T6SS-secreted effector TseG, interacts with the upstream chaperone TecG (DUF2169) and the downstream immunity protein TsiG (DUF3540). TseG exhibited antibacterial activity against *Escherichia coli* and enhanced virulence in maize, potato, and onion plants compared to a *tseG* deletion mutant [17]. Given the high conservation of this toxin-immunity pair across the *Pantoea* genus [17], *P. allii* may employ TseG in a similar functional role. Interestingly, several RHS domain-containing genes in *P. allii* strains DOAB1035 and MAI6007 appear to be pseudogenized, potentially due to the loss of their corresponding immunity genes. Additionally, numerous hypothetical protein-encoding genes found on *vgrG*-2 island may represent orphan immunity genes. While the VgrG effector of *P. agglomerans* pv. betae strain 4188 carries a C-terminal glucosaminidase domain targeting peptidoglycan [16], *P. allii* VgrGs lack this feature, suggesting a distinct mechanism of antibacterial activity to that of *P. agglomerans*. Instead, *P. allii* T6SS may rely on RHS domain-containing effectors such as the pentapeptide repeat domain-containing protein (COG1357) or uncharacterized hypothetical proteins within its *vgrG* islands to mediate bacterial competition and ecological adaptation,

In addition to the Type 6 Secretion System (T6SS), *P. allii* harbors Pantailocin islands (Fig. 5) resembling those found in *P. stewartii* subsp. *indologenes* ICMP10132 or *P. ananatis* ATCC35400 [20]. Pantailocins are R-type bacteriocins, bacteriophage-derived protein complexes that resemble contractile phage tails that selectively kill closely related *Pantoea* species [20]. The presence of both types of Pantailocin islands in *P. allii* suggests independent acquisition events rather than inheritance from a common ancestor. Populations of *P. allii* carrying both types of Pantailocins islands may gain a competitive advantage by eliminating Pantailocin-susceptible *P. ananatis* populations. While the production and killing activity Pantailocins found in *P. allii* remains to be investigated, high synteny and sequence homology of these islands to those in *P. stewartii* subsp. *indologenes* ICMP10132 or *P. ananatis* ATCC35400 suggest similar target specificity. It would be also interesting to determine whether the expanded *P. ananatis*-like Pantailocin island found in DOAB1051 encodes tailocins with altered target specificity.

## Conclusion

This study provided a valuable genomic, phylogenetic, and phenotypic characterization of the infrequently isolated *P. allii*, revealing distinct evolutionary lineages and diverse genetic loci relevant to its ecology, pathogenicity, and potential as a biocontrol agent. The detection of ice nucleation activity and HiVir gene cluster in strains isolated from environmental reservoirs such rainwater suggests possible dissemination routes of *P. allii* as an inoculum source for agriculture. While HiVir and Halophos are recognized onion virulence factors, their specific *in planta* targets still remain unknown and require further investigation. The diversity of T6SS-associated toxins and Pantailocin islands points to a complex network of interbacterial interactions that may support both competitive and pathogenic strategies of *P. allii*. Investigating the functional roles of these loci would enhance our understanding of *P. allii*’s persistence in plant-associated environments and their interaction with other microbes.

## Funding information

This work is supported by Specialty Crops Research Initiative Award 2019-51181-30013 from recommendations expressed in this publication are those of the author(s) and do not necessarily reflect the view of the U.S. Department of Agriculture.

The U.S. Department of Agriculture (USDA) is an equal opportunity provider and employer. Mention of trade names or commercial products in this publication is solely for the purpose of providing specific information and does not imply recommendation or endorsement by the USDA.

This work is supported by CSIC Grupos de Investigación I+D, Universidad de la República (Uruguay).

## Acknowledgements

We thank Tim Dumonceaux for making strain OXWO6B1 available for phenotyping.

The Program for the Development of Fundamental Science (PEDECIBA, Uruguay), postgraduate fellowship for S. De Armas from ANII-FNB (Uruguay).

Contribution of J.T. Tambong funded through Agriculture and Agri-Food Canada project# J-002272.

## Supplementary figures and tables

**Figure S1.**
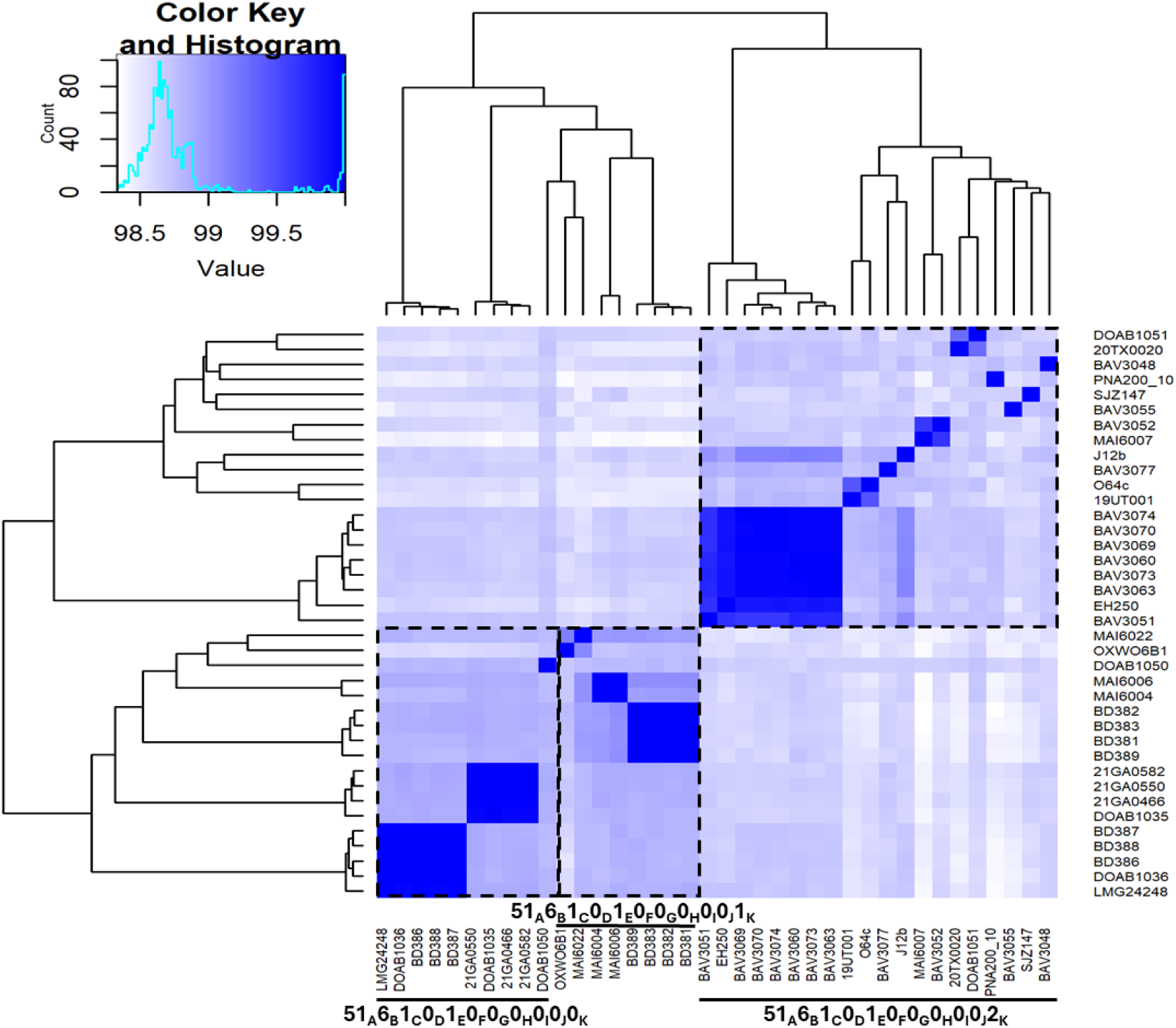
Heatmap of pairwise average nucleotide identity (ANI) between *P. allii* strains calculated by FastANI [38]. Lineages based on LINgroup are demarcated by dotted blocks.

**Figure S2.**
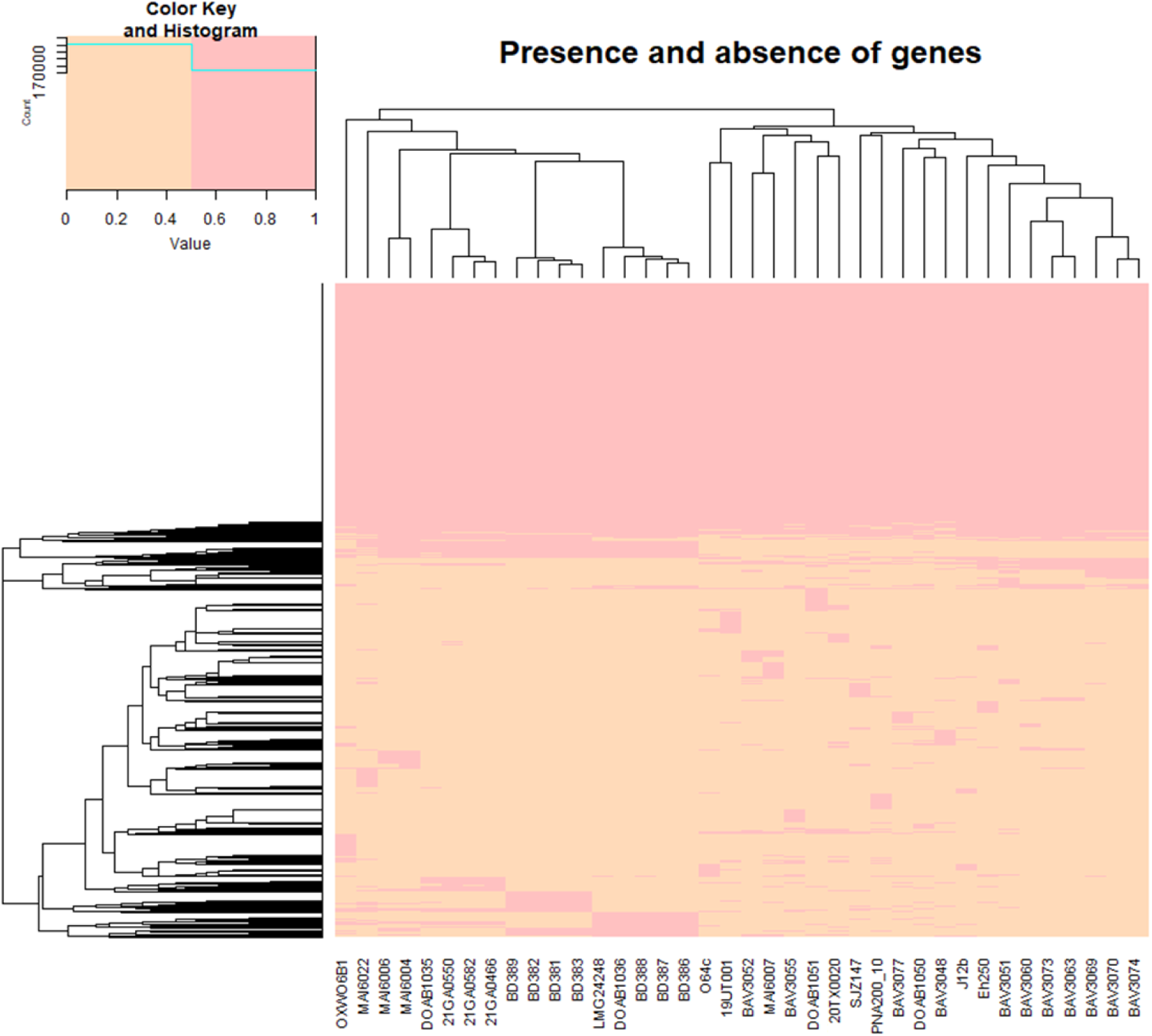
Presence and absence of genes dendrogram of 38 *P. allii* strains.

**Table S1.** Presence and absence (binary) data of virulence and secondary metabolite biosynthetic gene clusters (BGCs) detected in 38 *P. allii* strains.

**Table S2.** BUSCO completeness of *P. allii* genomes used in this study.

**Table S3**. Digital DNA-DNA hybridization values to the type *P. allii* LMG24248^T^ genome.

**Table S4.** QUAST genome assembly statistics of *P. allii* genomes used in this study.

**Table S5.** LINgroup assignment of 38 *P. allii* genomes by LINbase.

**Table S6.** Secondary metabolite biosynthetic gene clusters (BGCs) identified in 38 *P. allii* genomes by antiSMASH.

**Table S7.** Type secretion system identification by TXSS (Beav) in 38 *P. allii* genomes

**Table S8.** Identification of conjugative elements by CONJScan (MacSyFinder) in 38 *P. allii* genomes.

**Table S9**. Conserved domain search of vgrG-islands of *P. allii* T6SS-I.

**Table S10.** Conserved domain search of vgrG-island of *P. allii* T6SS-II.

**Table S11**. Phage island identification in 38 *P. allii* genomes by PHASTEST.

